# Unbounded Cell Growth and Proteome Imbalance in Doxorubicin-Induced Senescent RPE-1 Cells

**DOI:** 10.1101/2025.10.27.684969

**Authors:** Xili Liu, Matthew Sonnett, Marc W. Kirschner

## Abstract

Cellular senescence is traditionally viewed as a terminal state of cell-cycle arrest accompanied by widespread molecular remodeling, yet its underlying regulatory logic and progression remain poorly understood. Here, we combined quantitative phase microscopy and normalized Raman imaging with quantitative proteomic and phosphoproteomic profiling to examine human RPE1 cells undergoing doxorubicin-induced senescence. Senescent cells did not reach a steady state but instead exhibited sustained, unbounded growth over a 12 day period, marked by a continuous rise in dry mass and volume coupled with declining mass density. Time resolved proteomics revealed extensive and asynchronous remodeling across organelles, with lysosomal, ER, and Golgi proteins increasing in abundance, whereas nuclear and mitochondrial proteins declined, indicating large scale reorganization of cellular composition. Phosphoproteomic inference linked these structural shifts to regulatory signaling, confirming the expected downregulation of CDK activity while revealing coordinated activation of stress and DNA damage responsive kinases such as CAMK2D, DNAPK, and MARK family members. Together, these integrated data depict senescence as a dynamic, actively regulated state, maintained through coordinated remodeling of proteome composition and signaling activity rather than passive arrest. Our findings highlight how combining quantitative biophysical measurements with multi-layered molecular profiling exposes the regulatory architecture that sustains the senescent phenotype and its loss of internal homeostasis.

## Introduction

Cellular senescence has long been recognized as a state of stable cell cycle arrest accompanied by extensive molecular and functional changes(1). However, a precise and universally accepted definition remains elusive(2). Most efforts to characterize senescent cells rely on molecular markers such as p21, p16, senescence-associated-β-galactosidase (SA-β-gal) activity, and senescence-associated secretory phenotype (SASP) expression—but these markers vary from cell type to cell type, the nature of the inducing stimulus, and the timing of observation, limiting their generalizability(2). Although senescent cells were first discovered in cultured systems (3), the discovery of similar cellular features has been linked to aging, tissue regeneration, and chronic disease in animals (4), prompting growing interest in identifying them in vivo(5,6). However, the heterogeneous mixture of cell types and states in tissues makes this task even more challenging, as none of these molecular markers are exclusively specific to senescence(7). For instance, while p16 is commonly used to identify cell cycle–arrested cells, many terminally differentiated cells also express p16, (8). Thus, universal and robust markers for senescence are critical for advancing the field.

One widely observed feature across many senescent systems is cell size enlargement(9). Increased cell spreading and area expansion have been documented in senescent cells induced by various stimuli in vitro, including replicative stress, genotoxic agents, oncogene activation, viral infection, chemotherapeutics, and oxidative stress(10–15). It has been noted that size enlargement has also been observed in fibroblasts, epidermal keratinocytes, epithelial, and endothelial cells in aged animals(16–19), suggesting that size enlargement may be a general marker of senescence both in vitro and in vivo.

The size enlargement of senescence cells is particularly striking, given that cell size is tightly regulated in normal physiology(20). In proliferating cells, cell growth and the cell cycle are coupled to maintain a stable size distribution(21). For example, when the cell cycle is prolonged by PHA-848125, a CDK1/2 inhibitor, cell growth slows accordingly to preserve size at a similar level(22). In many settings, cell growth ceases when proliferation stops, such as in contact-inhibited monolayers or during terminal differentiation(23,24). Senescent cells, however, are notable for continuing to grow after permanent cell cycle arrest, resulting in cells of enlarged size. This raises several important questions. Do senescent cells reach a new steady-state size, or does excessive cell growth continue indefinitely? Size enlargement in senescent cells was usually measured by cell area or volume, which does not distinguish dilution from balanced mass increase (10–19). The question remains whether the observed geometrical size increase reflects real biomass accumulation or merely swelling? Mass density dilution has recently been proposed as a new feature of senescence cells(10,25–27). Do senescent cells represent a different homeostatic state of lower mass density, or does mass density continue to change over time? And ultimately, what do these changes in biophysical properties mean for senescent cell physiology?

To answer these questions, we characterized how key parameters of cell size, including dry mass, volume, and mass density, evolve during the process of senescence progression. To do this we need to employ newly developed high-precision biophysical measurement tools, which we have done in this report. To link such changes with underlying molecular processes, we performed sensitive quantitative proteomics and phosphoproteomics to dissect the molecular changes accompanying size enlargement. Finally, we consider what these changes may imply for the physiology and fitness of senescent cells.

Before turning to the results, we note that while senescence is most physiologically relevant in vivo, such as in aging and tissue repair, it remains technically difficult to perform accurate, high-resolution biophysical quantification and proteomic profiling for tissue samples. Therefore, we used a well-controlled in vitro model of senescence in cultured cells. Although this simplified 2D system carries limitations and may not fully recapitulate the complexity of tissue environments, we hope that discoveries made here will serve as basis for future in vivo investigations.

## Results

### Doxorubicin-Treated RPE-1 as a model for Cellular Senescence

RPE-1 cells treated with doxorubicin are a widely used model for studying cellular senescence in vitro(28,29). RPE-1 is a non-transformed, diploid human epithelial cell line with intact p53 and Rb pathways(30,31), making it suitable for investigating senescence in a genetically “wild-type” background. Doxorubicin, a chemotherapeutic agent that induces DNA double-strand breaks, triggers a robust DNA damage response and is commonly used to model genotoxic- and therapy-induced senescence(32). We chose this system to mimic a stressor that cells in aging organisms usually encounter, namely accumulated DNA damage(33). To minimize variability due to cell cycle re-entry, we maintained RPE-1 cells under continuous exposure to 100 nM doxorubicin and profiled them on Days 0, 1, 4, and 7 post-treatment.

To measure cell proliferation rate, we used EdU incorporation(6). EdU is a thymidine analog that is incorporated into DNA during replication and marks cells in the S phase(34). The assay revealed a dramatic drop in EdU-positive cells by Day 1, with negligible incorporation observed on Days 4 and 7, consistent with irreversible cell cycle arrest (Fig. 1A). The activity of the β-galactosidase measured at pH 6.0 is a widely used marker for cellular senescence, reflecting increased lysosomal mass(6,35). We observed a progressive increase in β-galactosidase activity, confirming that the cells had entered senescence (Fig. 1B).

**Figure 1.**
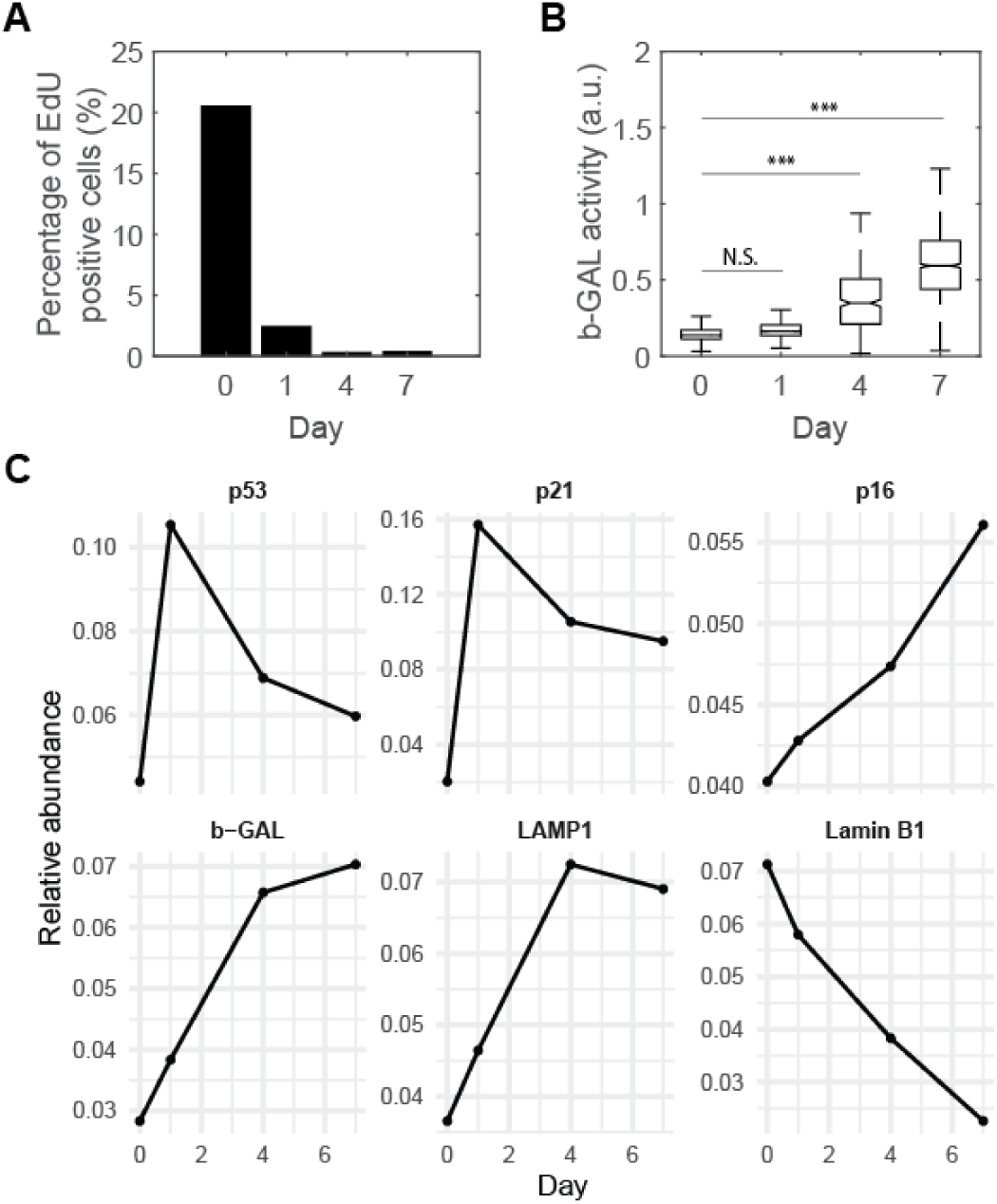
Doxorubicin-treated RPE-1 cells undergo cell cycle arrest and exhibit canonical markers of senescence. (A) Percentage of EdU-positive cells measured at Days 0, 1, 4, and 7 after continuous treatment with 100 nM doxorubicin. (B) Quantification of SA-β-galactosidase (β-GAL) activity. Box plots represent median, interquartile range, and 10th–90th percentiles. N.S. denotes no significant difference, while **** indicates p < 0.0001 by one-way ANOVA. (C) Relative abundance of canonical senescence markers from proteomic analysis.

Proteomic analysis captured dynamic changes in several canonical senescence markers (Fig. 1C). The level of p53 rose transiently, presumably in response to genotoxic stress caused by doxorubicin(36), and then declined as senescence became established. Among cell cycle inhibitors, p21 was rapidly upregulated as a direct transcriptional target of p53 but gradually declined over time, as its role in maintaining cell cycle arrest was overtaken by rising level of p16, a key enforcer of late senescence(37–40). In parallel, β-galactosidase and the lysosomal membrane protein LAMP1 increase in abundance, indicating elevated lysosomal content, while Lamin B1, a structural component of the nuclear lamina, decreased, reflecting nuclear morphological changes, both of which are consistent with known hallmarks of cellular senescence(6,7). Notably, the dramatic increase in β-galactosidase expression was consistent with the elevation of SA-β-gal activity assayed microscopically (Fig. 1B). Together, these data confirm that doxorubicin-treated RPE-1 cells underwent DNA damage- and p53-depedent cellular senescence(40), characterized by irreversible cell cycle arrest and increased lysosomal content.

### Senescent RPE-1 Cells Undergo Sustained, Unbounded Growth

To quantify cell growth during senescence, we measured cellular dry mass using Quantitative Phase Microscopy (QPM) (Fig. 2A) and mass densities using Normalized Raman Imaging (NoRI) (Fig. 2B), respectively. QPM is a label-free imaging technique that measures the phase shift, or optical path difference (OPD), caused by the refractive index difference between the cell and the surrounding medium(41). Because refractive index is proportional to solute concentration, integrating the OPD over the cell area enables quantification of cellular dry mass(42). We previously developed a computationally enhanced QPM (ceQPM) method that reduces measurement error to less than 2%(43). NoRI is a quantitative Stimulated Raman Scattering (SRS) microscopy method recently developed in our lab(27). It separates and normalizes signals from proteins, lipids, and water based on their characteristic Raman bands to determine their absolute concentrations, with an axial resolution of approximately 0.5 μm.

**Figure 2.**
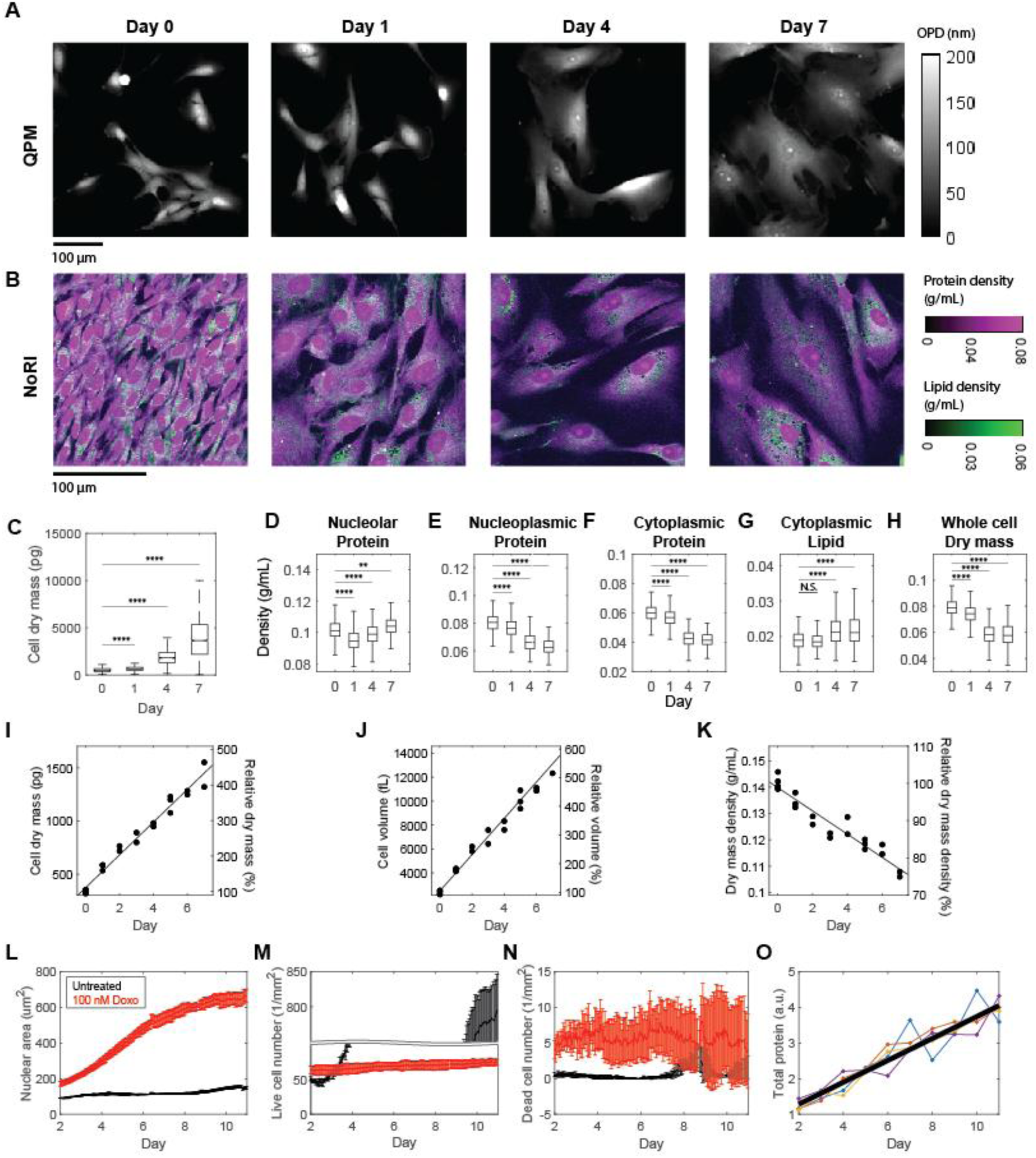
Senescent RPE-1 cells undergo sustained, unbounded growth. (A, B) Representative Quantitative Phase Microscopy (QPM) (A) and Normalized Raman Imaging (NoRI) (B) images of RPE-1 cells treated continuously with 100 nM doxorubicin, acquired at the indicated days. Scale bars, 100 µm. QPM grey scale indicates optical path difference (OPD); NoRI color scales indicate protein density (magenta) and lipid density (green). (C) Box plot of cell dry mass measured by QPM at Days 0, 1, 4, and 7. (D–H) Box plots of nucleolar (D), nucleoplasmic (E), and cytoplasmic (F) protein densities, cytoplasmic lipid density (G), and whole-cell dry mass density (H) measured by NoRI at Days 0, 1, 4, and 7. N.S., not significant; **p < 0.01; ****p < 0.0001 by one-way ANOVA. (I–K) Time course of median cell dry mass (I), volume (J), and dry mass density (K) measured daily from Day 0 to 7 in trypsinized single cells using QPM. Right axes show fold-change (%) relative to Day 0. Solid black lines indicate linear fits. (L–N) Average nuclear area (L), live cell number (M), and dead cell number (N) of control (Untreated, black) and 100 nM doxorubicin-treated (100nM Doxo, red) cells over time, tracked by IncuCyte using nuclear fluorescence. Error bars indicate standard deviation from three replicates. (O) Total protein abundance measured by BCA assay daily from Day 2 to Day 11. Each colored line represents an independent biological replicate; the black line shows the best linear fit across all replicates.

The results showed a continuous increase in cell dry mass from a mean of 541 to 4122 pg, a 7.6-fold increase from Day 0 to Day 7 (Table 1, Fig. 2C). While the nucleolar protein density remained within a small range between 0.095 and 0.104 g/mL (Table 1, Fig. 2D), nucleoplasmic protein density decreased by 22%, from 0.080 to 0.063 g/mL (Table 1, Fig. 2E), and cytoplasmic protein density declined by 30%, from 0.060 to 0.042 g/mL (Table 1, Fig. 2F). The drop in protein mass density was accompanied by a 17% increase in cytoplasmic lipid density, from 0.019 to 0.022 g/mL (Table 1, Fig. 2G). As a result, the overall cellular dry mass density decreased by 26%, from 0.079 to 0.058 g/mL, over the 7-day period (Table 1, Fig. 2H).

**Table 1.** Quantification of cell dry mass and mass densities during senescence progression. Mean, standard deviation (Std.), and coefficient of variation (CV, Std./Mean) of cell dry mass and compartment-specific mass densities measured at Days 0, 1, 4, and 7 in RPE-1 cells treated continuously with 100 nM doxorubicin. Dry mass was measured by QPM, and mass densities were measured by NoRI.

| Properties |  | Day 0 | Day 1 | Day 4 | Day 7 |
| --- | --- | --- | --- | --- | --- |
| Dry mass (pg) | Mean | 541.94 | 668.63 | 1866.50 | 4122.10 |
|  | Std. | 260.14 | 251.73 | 818.24 | 4129.40 |
|  | CV | 0.4800 | 0.3765 | 0.4384 | 1.0018 |
| Nucleolar protein density (g/mL) | Mean | 0.1017 | 0.0950 | 0.0985 | 0.1040 |
|  | Std. | 0.0062 | 0.0070 | 0.0072 | 0.0072 |
|  | CV | 0.0610 | 0.0737 | 0.0731 | 0.0692 |
| Nucleoplasmic protein density (g/mL) | Mean | 0.0801 | 0.0763 | 0.0664 | 0.0628 |
|  | Std. | 0.0068 | 0.0074 | 0.0067 | 0.0063 |
|  | CV | 0.0849 | 0.0970 | 0.1009 | 0.1003 |
| Cytoplasmic protein density (g/mL) | Mean | 0.0597 | 0.0566 | 0.0427 | 0.0416 |
|  | Std. | 0.0058 | 0.0061 | 0.0063 | 0.0065 |
|  | CV | 0.0972 | 0.1078 | 0.1475 | 0.1563 |
| Cytoplasmic lipid density (g/mL) | Mean | 0.0190 | 0.0188 | 0.0219 | 0.0223 |
|  | Std. | 0.0028 | 0.0023 | 0.0045 | 0.0054 |
|  | CV | 0.1474 | 0.1223 | 0.2055 | 0.2422 |
| Whole cell dry mass density (g/mL) | Mean | 0.0789 | 0.0741 | 0.0586 | 0.0583 |
|  | Std. | 0.0067 | 0.0073 | 0.0097 | 0.0107 |
|  | CV | 0.0849 | 0.0985 | 0.1655 | 0.1835 |

Despite the dramatic increase in dry mass and dilution in mass density, senescent cells became much more heterogeneous. We use the coefficient of variation (CV, defined as the standard deviation divided by the mean) to represent the variation within a population. The CV of dry mass increased 2.1-fold, from 48% on Day 0 to 100% on Day 7 (Table 1).

Previously, we reported that while cell mass doubles within one cell cycle, the mass density of proliferating cells is tightly regulated within a narrow range(26). This was supported by the low CVs observed on Day 0 for cytoplasmic protein density (9.7%) and whole-cell dry mass density (8.5%) (Table 1). By Day 7, however, these CVs had increased substantially, reaching 16% and 18%, respectively (Table 1). The CV of cytoplasmic lipid density also rose from 15% to 24% during this period (Table 1). All of these reflect increased cell-to-cell variability in senescent populations.

To assess growth dynamics during senescence progression at higher temporal resolution, we performed daily QPM measurements. QPM background subtraction requires a sparse cell distribution, which is difficult to maintain over extended time courses as cells grow and spread. To address this, we used trypsinized cells for measurement. In addition to dry mass, we estimated cell volume based on the area of rounded cells and calculated dry mass density by dividing dry mass by volume. Notably, volume measurement using the Coulter Counter is unreliable for senescent cells, as their swollen morphology makes them prone to lysis while passing through the measurement aperture. The area of rounded cells provides a more robust approximation of volume in this context. We found that both dry mass and volume increase approximately linearly during this period (46% per day in mass; 63% per day in volume) (Fig. 2I, J). Since dry mass increased more slowly than volume, dry mass density declined steadily by 3% per day (Fig. 2K). Unfortunately, for such measurements, senescent cells became increasingly resistant to trypsinization and fragile upon detachment; we therefore terminated these measurements on Day 7.

To circumvent the issue of cell lysis during detachment, we extended our observation period by using nuclear area and total protein abundance as proxies for cell size. We tracked cells expressing mCherry fused to nuclear localization sequences on both termini as a reporter for cell size(22), and recorded time-lapse movies using the IncuCyte live imaging system from Day 2 to Day 11. We found that average nuclear area continued to increase throughout the 9-day period, with a slower rate after Day 7, likely due to contact inhibition limiting cell spreading (Fig. 2L). Despite less than 25% increase in cell number over the entire period (Fig. 2M, N), total protein abundance increased linearly at a rate of ∼26% per day, as assayed by BCA (Fig. 2O), supporting the notion of continuous biomass accumulation.

These results demonstrate that senescent RPE-1 cells undergo unbounded growth, sustaining mass accumulation and volume expansion over an extended period without signs of plateau, accompanied by continuous mass density dilution.

### Proteomic Profiling Reveals Coordinated Remodeling of Cellular Architecture

To dissect the molecular basis of the sustained mass accumulation observed by QPM and NoRI imaging, we performed quantitative proteomic profiling of RPE-1 cells undergoing doxorubicin-induced senescence. The goal was to determine how global protein composition and subcellular organization evolve as cells continue to grow after cell-cycle arrest. Across a 12-day time course of continuous 100 nM doxorubicin treatment, we quantified approximately 5,000 proteins in four independent biological replicates, providing a detailed view of proteome remodeling during the transition from early to late senescence.

The resulting dataset showed exceptionally high reproducibility. PSMA3 was stable and exhibited nearly identical trajectories across all replicates (Fig 3A). In contrast, senescence markers such as β-galactosidase were strongly induced, confirming robust and synchronous senescence entry, with similarly strong biological agreement. Further underscoring the quality of the measurements, members of the same macromolecular complex displayed highly correlated behavior across replicates. For example, four of the MCM complex subunits measured declined in parallel, each showing a similarly steep decrease in abundance (Fig. 3B). This agreement within replicates and across functionally related proteins establishes that the dataset captures true biological trajectories with high fidelity.

**Figure 3.**
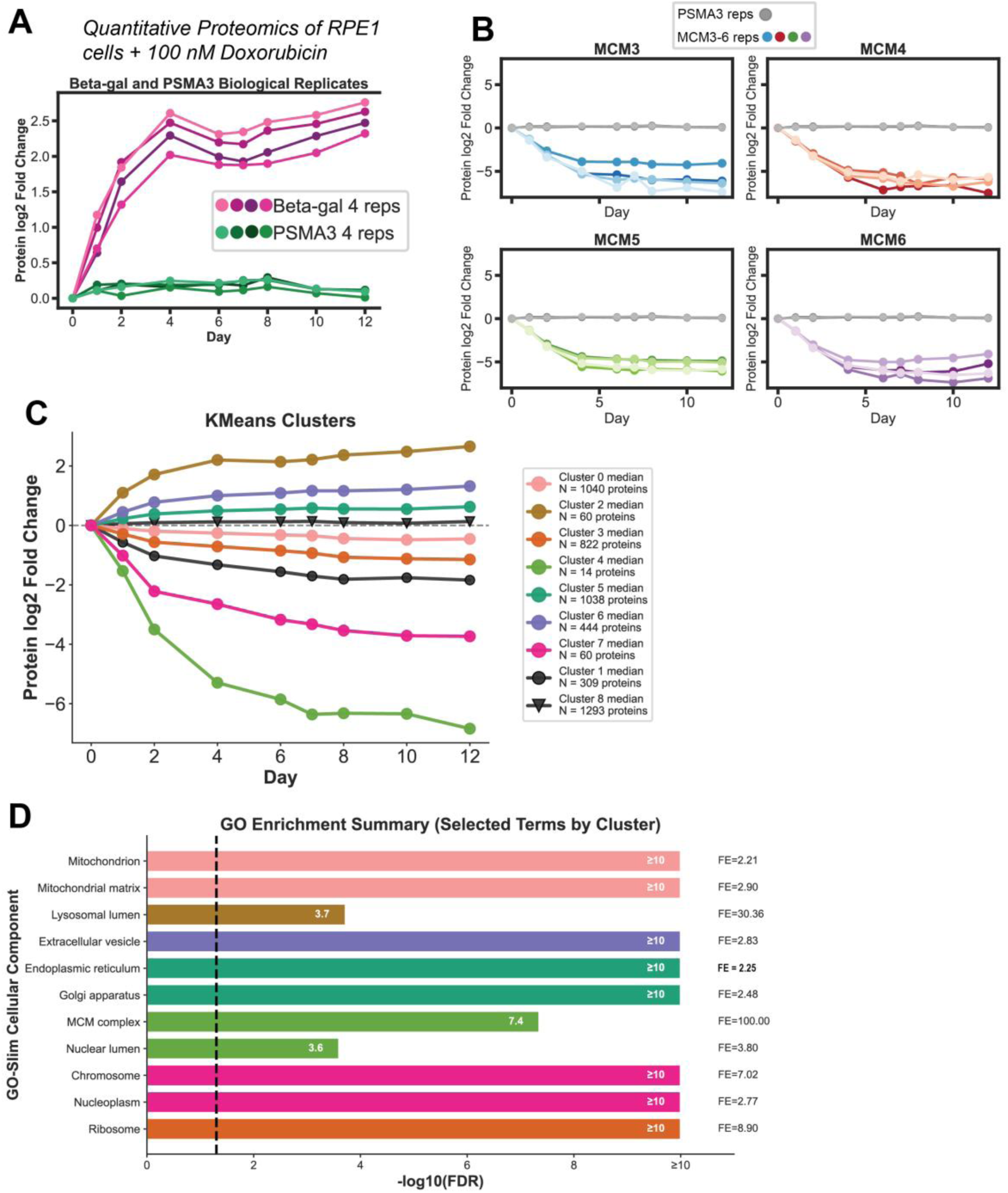
Quantitative proteomic profiling of RPE-1 cells treated with doxorubicin. (A) Time-course of protein abundance changes for β-galactosidase (magenta) and PSMA3 (green) biological replicates during 12 days of 100 nM doxorubicin treatment in the proteomics data. (B) Abundance trajectories for representative DNA-replication factors (MCM3–MCM6); each trace shows individual biological replicates with PSMA3 replicates shown in gray for comparison. (C) K-means clustering of all quantified proteins reveals seven distinct temporal expression patterns across the treatment course. (D) Gene Ontology (GO) enrichment analysis of selected representative cellular-component terms from each cluster. Bar colors are the same as the clusters in C. Bars indicate the significance of enrichment (x-axis: –log₁₀ FDR), with numbers on the right denoting the corresponding fold enrichment (FE).

Unsupervised K-means clustering of temporal trajectories separated the ∼5,000 quantified proteins into nine kinetic clusters (Fig. 3C). Rather than a single monotonic pattern, these clusters capture a spectrum of coordinated programs that evolve over the 12-day period. Some groups exhibit rapid and irreversible decline, others plateau, and a subset rise at different slopes as senescence deepens. Gene Ontology enrichment (Fig. 3D) links these temporal profiles to specific biological systems, revealing a coherent but highly asymmetric restructuring of the proteome. Clusters 4 (light green) and 7 (dark pink) are enriched for nucleoplasm, chromosomal, and nuclear-lamina proteins and dropped earliest and most sharply, marking an abrupt exit from the cell cycle. Ribosomal and RNA-processing factors (Cluster 3, Orange) followed a slower downward trajectory, indicating progressive suppression of translation and biosynthetic capacity. In contrast, proteins associated with lysosomes (Cluster 2, Brown), ER (Cluster 5, dark green), and the Golgi apparatus (Cluster 5, dark green)) accumulated over time, mirroring the gradual expansion of membranous compartments observed by imaging. Mitochondrial proteins (Cluster 0, light pink) largely declined, while select subsets, particularly redox and stress-related enzymes, remained stable or modestly increased.

### Organelle-level remodeling reveals disproportionate expansion of membranous compartments

Grouping proteins by organelle annotations highlighted heterogeneous scaling within each compartment (Fig. 4A–F). While some compartments expand in step with cell growth, others contract or reorganize, revealing that senescence represents a selective reallocation of cellular investment within each organelle.

**Figure 4.**
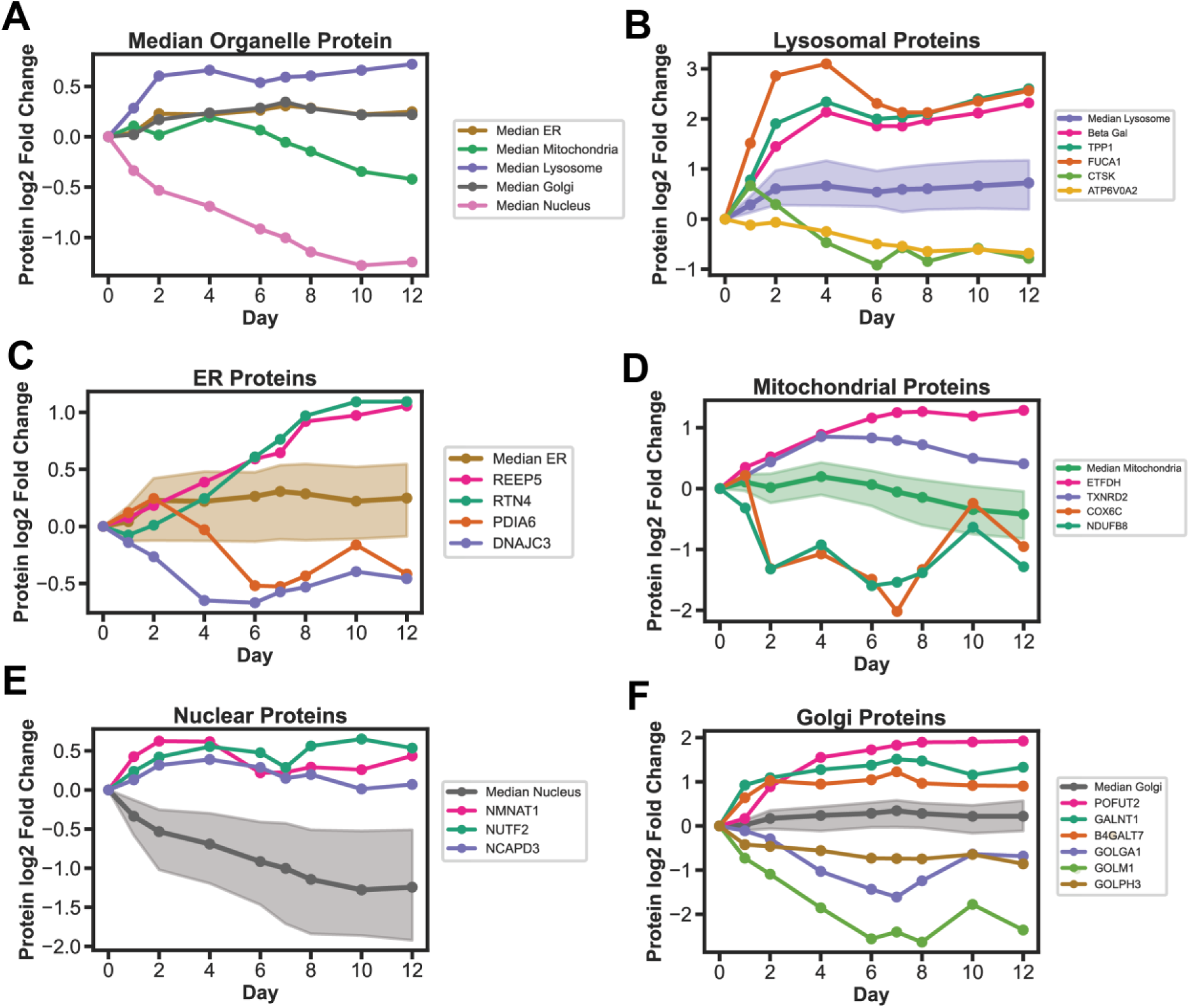
Organelle- and metabolism-specific protein scaling during senescence induction in RPE-1 Doxorubicin treated cells. (A) Median protein abundance trajectories for five major organelles, showing distinct scaling behaviors over 12 days. (B–F) Examples of differentially scaling proteins and median ± IQR traces for each compartment: (B) lysosome, (C) ER, (D) mitochondria, (E) nucleus, and (F) Golgi. Protein trajectories are plotted as individual proteins with median ± IQR curves.

Lysosomes illustrate this principle most clearly. The median abundance of lysosomal proteins increases ∼1.5-fold, yet individual enzymes behave very differently (Fig. 4B). Canonical hydrolases such as β-galactosidase, whose activity defines the SA-β-gal senescence marker (6, 35) along with TPP1 and FUCA1, rise sharply, suggesting enhanced catabolic capacity. Others move in the opposite direction: CTSK declines and the proton-pump subunit ATP6V0A2 falls (44), implying that acidification and substrate specificity are being re-tuned rather than uniformly amplified. Even within one organelle, the proteome separates into expanding and contracting modules, reflecting functional specialization of the lysosomal system as cells enlarge.

The endoplasmic reticulum follows a related but distinct pattern (Fig. 4C). Shape-stabilizing proteins RTN4 and REEP5 which are architects of the ER’s tubular network (45) show some of the strongest increases, consistent with a remodeling of membrane geometry to accommodate greater surface area and trafficking demand. By contrast, the folding regulators PDIA6 and DNAJC3 decline. PDIA6 catalyzes disulfide-bond isomerization, and DNAJC3 inhibits the PERK arm of the stress response; their reduction suggests that quality-control capacity does not scale proportionally with membrane expansion. Thus, the ER may become physically larger but compositionally rebalanced, with more scaffolding for export and relatively less investment in stress-buffering enzymes.

The Golgi apparatus mirrors the ER’s modest upward trend but with even greater internal contrast (Fig. 4A,F). The glycosylation enzymes POFUT2, GALNT1, and B4GALT7 increase prominently, consistent with elevated glycan-processing throughput, whereas structural and tethering factors such as GOLGA1, GOLM1, and GOLPH3 decline. These opposing movements indicate that the Golgi’s biosynthetic machinery scales differently from its scaffolding and transport framework, emphasizing selective adjustment of glycosylation activity rather than wholesale enlargement of the organelle.

Mitochondria tell the opposite story. Their median protein abundance drops steadily after a short plateau (Fig. 4A, D). Redox-associated enzymes like ETFDH and TXNRD2 are modestly elevated, while respiratory-chain components COX6C and NDUFB8 fall sharply. This divergence implies that antioxidant defenses persist even as oxidative phosphorylation capacity wanes, consistent with a gradual functional down-shift of mitochondrial metabolism.

The nuclear proteome shows the most pervasive contraction (Fig. 4E). Structural and chromatin-associated proteins decline throughout the time course, matching the nuclear shrinkage and density loss measured by NoRI (29). However, multiple proteins such as NMNAT1, NUTF2, and NCAPD3 remain stable or increase slightly, maintaining NAD⁺ metabolism, nucleo-cytoplasmic transport, and chromatin organization despite the broader reduction in DNA-replication and transcriptional machinery.

Across all compartments, these patterns converge on a common theme: senescence remodels the cell unevenly. Lysosomes, ER, and Golgi expand or reconfigure their internal composition, while mitochondria and the nucleus contract. When integrated with QPM and NoRI measurements showing sustained mass accumulation and cytoplasmic protein dilution, the proteomic data indicate that cellular growth continues, but different subsystems change at different rates. The result is an increasingly unbalanced proteome with an expansion of membranes and trafficking machinery that is not matched by proportional adjustments in biosynthetic, nuclear, or mitochondrial components.

### Kinase activity landscapes reveal shared and divergent signaling responses to Doxorubicin and Palbociclib

The large-scale but uncoordinated proteome remodeling observed across organelles suggests that at least some proteome changes during senescence may be actively regulated rather than a passive consequence of cell growth. Coordinating the suppression of nuclear and biosynthetic programs with the expansion of secretory and degradative compartments requires dynamic control by intracellular signaling networks. Because many of these pathways act through protein phosphorylation, a rapid and reversible mechanism for integrating stress, metabolic, and growth cues, we next examined how phosphorylation patterns change during senescence.

To connect phosphorylation events to their upstream regulators, we used KINAID (66), an orthology-based framework that infers relative kinase activity from large-scale phosphoproteomic data. KINAID integrates experimentally derived kinase–substrate specificity scores with measured phosphosite intensities to estimate activity changes for ∼380 of kinases simultaneously, allowing phosphorylation data to be interpreted in terms of pathway activity rather than individual sites.

To determine whether the resulting signaling programs were specific to one inducer or reflected a shared regulatory architecture, we extended our analysis beyond doxorubicin-treated RPE-1 cells. We compared two mechanistically distinct routes to cell-cycle exit: DNA-damage–induced senescence triggered by Doxorubicin and CDK4/6 inhibition– induced senescence caused by Palbociclib, in both epithelial RPE-1 and fibroblast IMR90 cells. Quantitative phosphoproteomics time courses across all four conditions identified roughly 25,000 unique phosphosites. This design enabled comparison of signaling networks that define senescence induced by genotoxic stress versus CDK inhibition, and across distinct cellular contexts capable of stable cell-cycle withdrawal.

Kinase–substrate enrichment analysis revealed broad remodeling of phosphorylation-based signaling across both cell types and perturbations. Volcano plots for each condition (Fig. 5B–E′) illustrate these activity shifts, with numerous kinases showing significant changes in both Doxorubicin- and Palbociclib-treated cells. Despite differences in magnitude, all four datasets shared a dominant trend of CDK and CLK suppression alongside activation of selected stress-response kinases, motivating an integrated comparison of signaling architecture across treatments and cell types.

**Figure 5.**
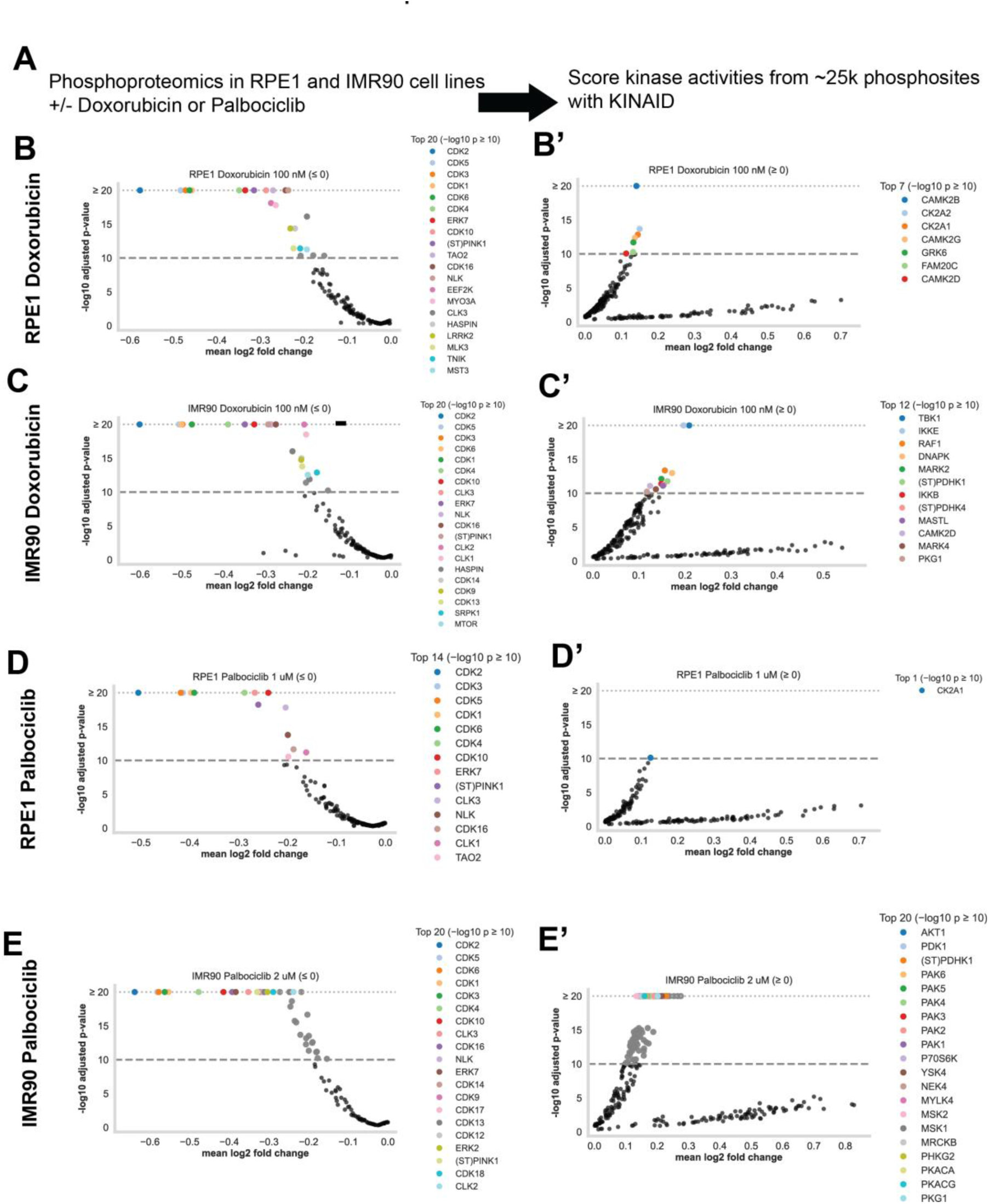
Kinase activity profiles inferred from phosphoproteomics in RPE-1 and IMR90 cells treated with Doxorubicin or Palbociclib. **(A)** Overview of the experimental and computational workflow. Phosphoproteomic datasets from RPE-1 and IMR90 cells treated with Doxorubicin or Palbociclib were analyzed with KINAID to infer relative kinase activity from ∼25,000 quantified phosphosites. **(B–E)** Volcano plots showing kinases with altered activity under each treatment condition (|mean log₂ fold change| ≥ 0.1, −log₁₀ adjusted p ≥ 10). Left panels (B–E) show kinases with decreased activity (mean ≤ 0); right panels (B′–E′) show kinases with increased activity (mean ≥ 0). Colored points denote the top significantly regulated kinases in each condition, with representative examples labeled in the accompanying legends.

The most prominent and consistent feature across conditions was a coordinated reduction in activity of cell-cycle and RNA-processing kinases (Fig. 6A–C). This down-regulated module included the CDK family (CDK1/2/3/4/5/6/10/16) and CLK1/3, together with mitotic regulators HASPIN and NLK, marking a broad shutdown of proliferative and chromatin-associated signaling programs. Additional kinases such as ERK7 and PINK1 also declined, extending this suppressed core into stress- and organelle-linked pathways. The effect was most pronounced in Palbociclib-treated samples, where CDK and CLK repression dominated the landscape, while Doxorubicin-treated IMR90 cells displayed the same repressed core accompanied by a broader stress-response signature.

**Figure 6.**
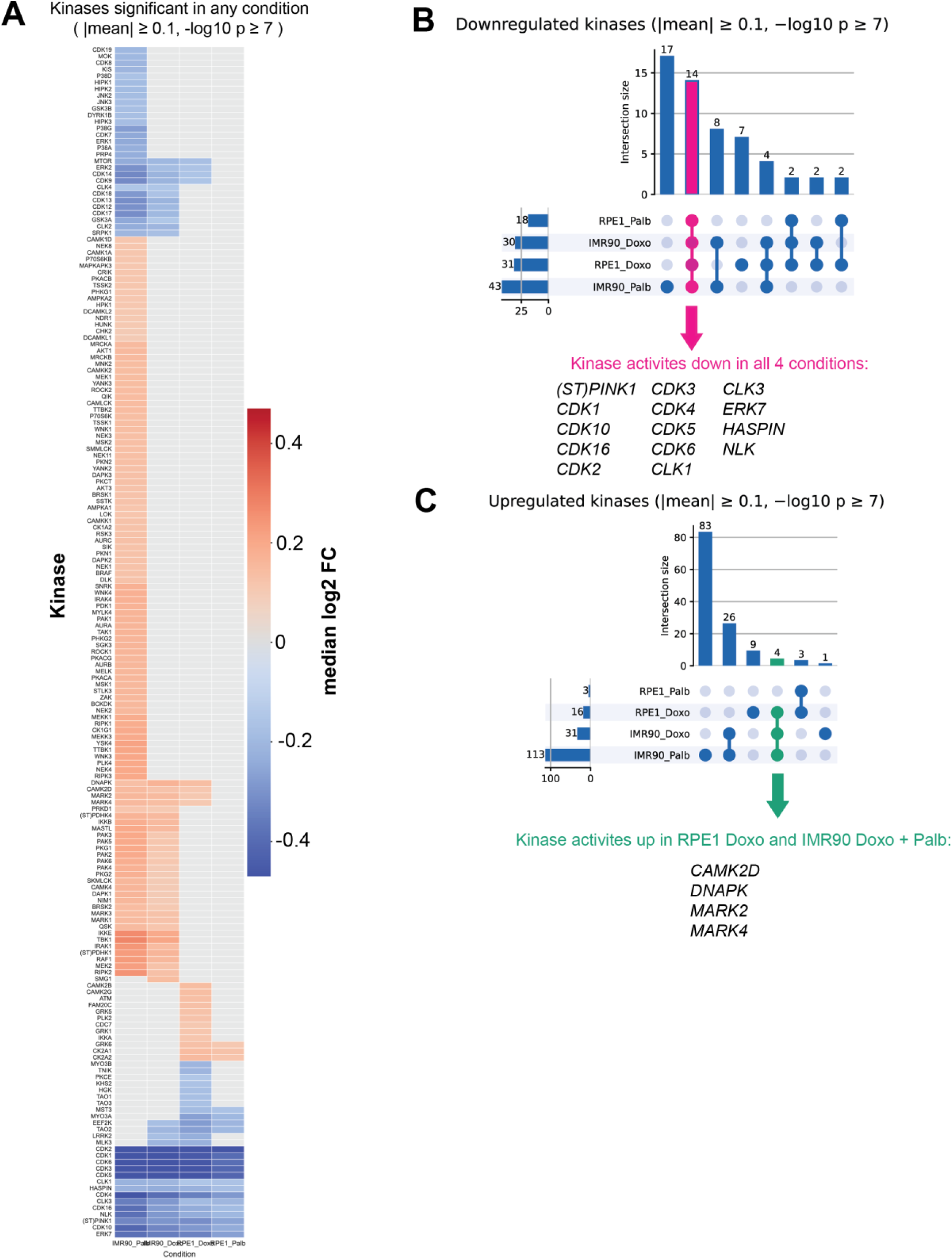
Comparative kinase activity responses to Doxorubicin and Palbociclib in RPE-1 and IMR90 cells. **(A)** Heatmap showing kinases significantly altered in at least one condition (|mean log₂ fold change| ≥ 0.1, −log₁₀ adjusted p ≥ 7) following Doxorubicin or Palbociclib treatment in RPE-1 or IMR90 cells. Red indicates increased and blue decreased inferred kinase activity relative to control; gray denotes non-significant changes. **(B)** UpSet plot of downregulated kinases across the four conditions. Fourteen kinases including multiple CDKs, CLKs, ERK7, NLK, and HASPIN were consistently decreased. **(C)** UpSet plot of upregulated kinases. Four kinases (CAMK2D, DNAPK, MARK2, MARK4) were commonly increased in RPE-1 Doxorubicin and both IMR90 Doxorubicin and Palbociclib treatments.

In contrast, several kinases associated with stress adaptation and cytoskeletal regulation were activated, including CAMK2D, DNAPK (PRKDC), MARK2, and MARK4 (Fig. 6C). These kinases integrate calcium, metabolic, and DNA-damage signals, indicating that both genotoxic and CDK-inhibitory stress converge on a shared signaling framework defined by CDK/CLK repression coupled to stress-kinase activation.

Collectively, the proteomic and phosphoproteomic results portray senescence as a regulated and actively maintained state rather than a terminal arrest. A conserved network of CDK and RNA-processing kinases is repressed, while stress- and repair-associated kinases remain engaged, coordinating the extensive remodeling of organelles and biosynthetic pathways. These findings establish a signaling blueprint for senescence and provide a foundation for exploring how distinct kinase programs sustain long-term growth arrest and stress tolerance.

## Discussion

### A Dynamic View of Senescence

In this study, we quantitatively tracked the biophysical and molecular remodeling of RPE-1 cells undergoing doxorubicin-induced senescence. By combining high-precision measurements of cell dry mass, volume, and cell mass density, coupled with high-resolution proteomic and phosphoproteomic profiling, we demonstrated that senescent cells exhibit sustained, unbounded growth over time, accompanied by extensive proteome remodeling and activation of stress response pathways. Rather than entering a stable end state, senescent cells continue to evolve, adapting their metabolism and stress response to maintain viability despite internal imbalance. The extended time course and high temporal resolution of sampling in these experiments allowed us to capture much more dynamic changes in both cell size and molecular composition than previously reported(29,54), offering a broad perspective on senescence progression. Principal component analysis further supports the view that senescence may proceed through multiple stages, rather than a single binary transition.

Beyond documenting growth itself, the proteomic data provide a molecular view of how senescent cells reorganize their contents as they enlarge. The time-resolved proteome revealed asynchronous remodeling across organelles with lysosomal and secretory proteins gradually accumulating, while nuclear and mitochondrial components decline. This suggested that senescent growth is accompanied by structural re-allocation rather than uniform biosynthetic scaling. Phosphoproteomic inference further connected these compositional changes to signaling control, showing a conserved repression of CDK and RNA-processing kinases coupled to activation of stress- and DNA-damage–responsive pathways. Together, these molecular layers link the biophysical growth phenotype to its upstream regulatory circuitry, emphasizing that senescence is an actively maintained and continuously remodeled state.

### Rethinking the Definition of Senescence

Whether senescence constitutes a specific cell state or a collection of diverse states sharing some common features remains an open question. Senescent cells induced by different triggers and originating from different cell types often exhibit only a subset of the “senescence markers”(7), and they differ in their SASP expression profiles(54). Moreover, senescence can have either beneficial or detrimental effects depending on the biological context(55). This variability complicates efforts to define senescence as a single state and may in part explain the current lack of consensus surrounding its molecular composition and circuitry. The view that senescence is a temporal progression rather than a discrete final state might help us reconcile some of these inconsistencies in the interpretation of cellular senescence. Characterizing molecular trajectories with higher time resolution through senescence progression may be more productive than searching or a fixed set of markers to define the senescence state.

### Physiological Hallmarks of Senescence

One key question is what features are most fundamental and universal to senescent cells. Irreversible or permanent cell cycle arrest is often used as the defining characteristic(1), yet this criterion has limitations. Irreversibility is a relative concept that depends on the observation window; notably several studies have shown that escape from senescence is a common feature of cultured cells(40,56). Moreover, a definition of senescence based on arrest of the cell cycle from a proliferative state implies that only proliferative cells are capable of becoming senescent. However, certain terminally differentiated, non-proliferative cells (e.g., neurons) can exhibit senescence-like phenotypes(9,57), including expression of SA-β-gal, p21, p16, and SASP(57). Additionally, even in proliferating cells, cell cycle arrest is not always required for cells to display features commonly associated with senescence. For instance, cells under chronic stress can undergo size enlargement without evident cell cycle extension(58).

Our observation of unbounded growth in vitro, marked by continuous increases in dry mass and volume, along with a decrease in mass density, raises the possibility that loss of internal homeostasis may drive cells into a pathological state. In this state, cells lose their original identity and function yet remain viable and continue to grow. Such behavior could represent a consistent and intrinsic aspect of senescence physiology and may be worth testing across diverse senescence models in vivo.

It is important to note that while we observed mass density dilution alongside size enlargement, this phenomenon may not directly translate to in vivo settings. Mass density is influenced by mechanical, osmotic, and metabolic factors(24,59), all of which differ significantly between in vitro and tissue environments. Moreover, we observed opposing changes in protein and lipid densities during senescence, suggesting that total dry mass density may increase or decrease depending on the relative magnitude of these shifts. Given this complexity, we propose that increased lipid density, rather than overall cytoplasmic dilution, may serve as a more robust and characteristic marker of senescent cells in situ.

Together, these observations suggest that size enlargement, mass density alteration, and the breakdown of internal homeostasis may constitute core physiological hallmarks of senescence that complement existing molecular markers. Consistent with these physical changes, the proteomic data highlight a coordinated loss of nuclear and mitochondrial machinery and a compensatory rise in secretory pathways, mirroring the same imbalance at the molecular level.

### Implications for Aging

Physiological features such as size enlargement, mass density changes, and loss of internal homeostasis may be especially relevant for understanding the role of senescent cells in aging. Senescent cells have been implicated in chronic inflammation, tissue microenvironment remodeling, and tumor-promoting signaling through paracrine effects(60), suggesting that even a small number of such cells could have disproportionate impacts on tissue function. For these effects to contribute meaningfully to aging-related pathology, senescent cells must influence the physiology of a substantial number of neighboring cells. Therefore, beyond identifying individual senescent cells, it may be useful to assess broader deviations from homeostatic norms across cell populations—such as changes in size, density, and proteome composition—as indicators of tissue-level dysfunction. These physiological features may serve as meaningful readouts in efforts to understand aging processes and evaluate interventions aimed at extending healthspan.

### Parallels Between Senescence and Cancer

The senescence program shares notable features with certain aspects of cancer biology. Despite their cell cycle arrest, senescent cells resist apoptosis, undergo metabolic rewiring, and increase lipid accumulation, similar to cancer cells, which reprogram their metabolism to support growth and survival(61,62). These similarities raise the possibility that senescent and cancer cells may exploit overlapping mechanisms to evade cell death and adapt to stressful environments. Notably, senescent and cancer cells often coexist in aging tissues(39). A deeper understanding of their shared traits could reveal common vulnerabilities and inform therapeutic strategies aimed at selectively targeting both cell types.

### Limitations and Future Directions

This study has important limitations. Most of the work focused on a single epithelial cell line (RPE-1), a single senescence inducer (doxorubicin), and an in vitro environment that lacks the complexity of tissue context. Senescence in vivo is shaped by interactions with neighboring cells, immune components, and extracellular matrix(63), which are not captured in this model. Additionally, while we inferred functional consequences from proteomic remodeling, follow-up studies are needed to directly test the roles of individual pathways, especially those involved in metabolism and stress adaptation. The kinase-activity signatures identified here marked by sustained CDK suppression and activation of stress- and repair-associated kinases provide concrete hypotheses for how signaling networks maintain senescent cell viability and may be experimentally testable through targeted perturbations. While sustained CDK repression is hardly unexpected, the broader cohort of kinase activities that rise or fall during senescence remains far less understood and warrants systematic exploration to uncover new signaling mechanisms unique to this state.

Nevertheless, by bridging quantitative biophysical measurements with highly quantitative proteomic profiling, our study offers a new and comprehensive view of how senescent cells grow, survive, and remodel over time. It underscores the importance of considering senescence as a dynamic and heterogeneous process, and highlights the potential of using physical properties—such as cell size and mass density—as robust and functionally meaningful markers of senescent cells. Future work extending these analyses across cell types, senescence triggers, and in vivo models will be critical for refining our understanding of senescence and its role in health and disease.

## Materials and Methods

### Cell Culture and Senescence Induction

RPE-1 (CRL-4000) cells were purchased from the ATCC. RPE-1 Nu-mCherry cells were generated by lentiviral infection. Single clones with stable expression were selected. Cells were cultured at 37°C with 5% CO₂ in Dulbecco’s Modified Eagle’s Medium (DMEM; 11965, Thermo Fisher Scientific) supplemented with 10% fetal bovine serum (FBS; 16000044, Thermo Fisher Scientific), 1% penicillin/streptomycin (15140122, Thermo Fisher Scientific), 25 mM HEPES (15630080, Thermo Fisher Scientific), and 10 mM sodium pyruvate (11360070, Thermo Fisher Scientific). Senescence was induced using 100 nM doxorubicin (S1208, Selleckchem). During IncuCyte experiments, 50 nM YOYO-1 (Y3601, Thermo Fisher Scientific) was added to the medium to label dead cells.

For NoRI measurements, cells were seeded on 55 mm glass-bottom dishes with 30 mm microwell #1.5 coverglass (D60-30-1.5-N, Cellvis) and fixed with 4% paraformaldehyde in PBS (diluted from 8% stock, RT 157–8, Electron Microscopy Sciences) for 30 minutes before imaging. Other microscopy experiments were performed using 6-well glass-bottom plates (P06G-1.5-20-F, MatTek), except for trypsinized cells. Trypsinized samples were prepared by treating cells with 0.05% trypsin-EDTA (25300054, Thermo Fisher Scientific), resuspending them in medium, adding them to the center of a circular imaging spacer (13 mm diameter, 247462, Grace Bio-Labs) on a microscope slide, and covering them with a coverslip.

### EdU Incorporation Assay

EdU incorporation was measured using the Click-iT™ Plus EdU Cell Proliferation Kit for Imaging, Alexa Fluor™ 647 dye (C10640, Thermo Fisher Scientific). Cells were pulse labeled with 20 µM EdU (5-ethynyl-2-deoxyuridine) for 2 hours. Click-iT™ chemistry was performed according to the manufacturer’s instructions. After conjugation, cells were stained with 10 μM Hoechst 33342 (62249, Thermo Fisher Scientific) for 30 minutes and imaged using a fluorescence microscope at 10× magnification.

### SA-β-galactosidase Staining and Quantification

SA-beta-galactosidase activity was detected using the CellEvent™ Senescence Green Detection Kit (C10850, Thermo Fisher Scientific). Cells were fixed in 2% paraformaldehyde in PBS for 10 min. The assay was conducted per the manufacturer’s instructions. Cells were then stained with 10 μM Hoechst 33342 and 500 ng/mL Alexa Fluor™ 568 NHS Ester (A20003, Thermo Fisher Scientific) for 30 minutes, and then imaged using fluorescence microscopy at 10× magnification. SA-β-gal activity was quantified as the green-to-red fluorescence ratio within the entire cell(26).

### Fluorescence Microscopy

Fluorescence microscopy was conducted on a Nikon Eclipse Ti2 microscope equipped with the Perfect Focus System (PFS) and an ORCA-Fusion BT Digital CMOS camera (Hamamatsu Photonics, Japan). Images were acquired using Nikon NIS-Elements AR software version 5.42.06 with the High Content Analysis module.

### Quantitative Phase Microscopy (QPM)

QPM images were acquired using a SID4-sC8 camera (Phasics, France) based on quadriwave lateral shearing interferometry (QWLSI). A Plan Apo λ 10× N.A. 0.45 objective and an LWD N.A. 0.52 condenser were used with the aperture diaphragm minimized. A halogen lamp was used for transmitted light. Image processing and cell mass quantification were performed according to the computationally enhanced QPM (ceQPM) method previously developed in our laboratory(43).

### Normalized Raman Imaging (NoRI)

Protein and lipid densities were quantified from stimulated Raman scattering (SRS) images following previously published protocols(26,27). SRS images were acquired at 2853 cm^−1^,2935 cm^−1^, and 3420 cm^−1^ bands (corresponding to methylene, methyl, and water vibrational bands, respectively) using a custom-built spectral-focusing femtosecond SRS microscope. The three-channel SRS images were spectrally unmixed into protein, lipid, and water components using reference spectra from bovine serum albumin in water, dioleoyl-phosphocholine in per-deuterated methanol, water, and per-deuterated methanol. The unmixed images were converted to absolute concentrations by normalizing the sum of the three components at each pixel. Dry mass density was calculated as the sum of protein and lipid mass densities.

### IncuCyte Live-Cell Imaging and Analysis

Cells were seeded in 24-well clear TC-treated plates (3524; Corning) with three replicate wells per condition and 1 mL of medium per well. Media was refreshed every 2–3 days by replacing 0.5 mL. Time-lapse imaging was performed using an IncuCyte ZOOM system (Sartorius) with a 20× objective. Nuclei and dead cells were detected using nuclear fluorescence and YOYO-1 staining, respectively(64). Average nuclear area, live and dead cell number densities were measured using the IncuCyte’s built-in software.

### Image Processing and Statistical Analysis

Images from fluorescence microscopy, QPM, and NoRI were processed using custom MATLAB (MathWorks) or ImageJ (NIH) scripts(21,26). Segmentation was based on morphological operations, thresholding, and watershed algorithms, except for nuclear segmentation in NoRI images, which was carried out using Cellpose(65). Quantitative measurements and statistical analyses were performed using custom MATLAB scripts. One-way ANOVA was used for group comparisons. Statistical significance was denoted as follows: N.S., p > 0.05; * p < 0.05; ** p < 0.01; *** p < 0.001; **** p < 0.0001.

### Proteomics Sample Preparation

Proteins were reduced by adding 5 mM dithiothreitol (DTT) (1 M stock in water; Oakwood Chemical #3483-12-3) and incubating at 60 °C for 15 min. Samples were then sonicated on ice for 10 × 15 s bursts at an amplitude of 60, clarified by centrifugation at 20,000 × g for 15 min at 4 °C, and only a faint pellet was typically visible.

To alkylate cysteine residues, 50 mM iodoacetamide (1 M stock in anhydrous N,N-dimethylformamide; BioUltra, Millipore Sigma #I1149; DMF #227056) was added and samples were kept in the dark for 1 h. Alkylation was quenched by introducing 30 mM DTT.

Proteins were precipitated using a chloroform/methanol extraction as described previously (67). The resulting protein disc was resuspended to ∼ 2 µg µL⁻¹ in 6 M guanidine HCl / 10 mM EPPS, pH 8.5. For digestion, 50 µg of protein from each sample was diluted to 2 M GuHCl with 10 mM EPPS (pH 8.5) and incubated with 20 ng µL⁻¹ LysC (Wako) for 14 h. The mixture was further diluted to 0.5 M GuHCl, supplemented with an additional 20 ng µL⁻¹ LysC and 10 ng µL⁻¹ trypsin (sequencing-grade modified; Promega #V5111), and digested for 16 h at 37 °C.

Following digestion, solvent was removed in vacuo, and peptides were resuspended in 500 mM EPPS (pH 8.0) at ∼ 1 µg µL⁻¹. 50 µg of peptide was labeled with 20 µL of TMTpro reagent (Thermo #A44520; 20 µg µL⁻¹ in anhydrous acetonitrile) for 2 h at room temperature. The reaction was quenched with 10 µL of 5 % hydroxylamine (Millipore Sigma #438227). Labeled samples were pooled, dried in vacuo, acidified to pH ≤ 1 with trifluoroacetic acid (TFA), clarified at 20,000 × g for 10 min at 4 °C, and desalted on a Sep-Pak C18 cartridge (Waters). 5 µg of TMT labeled peptides were analyzed in each LC-MS/MS run.

### LC-MS

Mass-spectrometric analyses were performed on a Thermo Orbitrap Eclipse Tribrid operating in data-dependent acquisition mode. Survey scans covered m/z 300–1073 at a resolution of 120,000, with a 200 ms maximum injection time, AGC target 1 × 10⁶, and RF lens 60 %. The instrument was coupled to a Thermo EASY-nLC 1200 system. Each run lasted 4.5 h, using a gradient from 4% to 19 % solvent B (solvent A: 0.125 % FA, 2 % DMSO; solvent B: 0.125 % FA, 80 % ACN, 2 % DMSO).

Chromatography was performed on in-house-packed 100/360 µm i.d./o.d. fused-silica columns containing ∼ 0.5–1 cm of Magic C4 resin followed by 40 cm of 1.8 µm ReproSil-Pur C18 resin (76). A custom 60 °C column heater was used, and peptides were electrosprayed at 2.6 kV through a PEEK micro-tee. Four LC–MS experiments were acquired using FAIMS compensation voltages of −32, −42, −52, and −62 V, operated at standard resolution with a total carrier-gas flow of 3.5 L min⁻¹.

Dynamic exclusion was set to 60 s (±10 ppm). MS/MS precursors were isolated in the quadrupole with a 0.4 m/z window and fragmented by HCD (37 % 29). Fragment spectra were acquired in the Orbitrap at 60,000 resolution, from m/z 110 to 2000, using an AGC target of 5 × 10⁴ and a 120 ms maximum injection time. High m/z TMTproC reporter ions were quantified within a 20 ppm tolerance and corrected for isotopic impurities as described previously (68).

For phosphoproteomics experiments, enriched phosphopeptides were analyzed with the same instrument settings for 4.5 hour runs, but only a single CV (ranging from -35 to -70 by steps of 5) was used for each run. The low m/z TMT reporter ions were quantified (69).

### Proteomics Data Analysis

Spectra were searched essentially as described previously (70), substituting the Comet search engine for Sequest. The precursor tolerance was 50 ppm, with variable modifications of oxidation (Met +15.9949 Da) and deamidation (Asn +0.9840 Da). Static modifications included carbamidomethylation (Cys +57.0215 Da) and TMTpro (N-termini/Lys +304.2066 Da). False-discovery rates were controlled using a target–decoy approach to 1 % at the peptide level (71), and proteins were filtered to 1 % FDR using Protein Pickr (72). Phosphoproteomics spectra were searched with a variable phosphate modification (STY +79.9663 Da).

### Kinase-Substrate Prediction and Annotation

Kinase–substrate relationships were inferred using KINAID (66) with default parameters. According to the authors instructions, a motif input column for each phosphorylation site was constructed with the 5 n-terminal and 4 c-terminal amino acids relative to the site quantified in the mass spectrometry data. This was provided along with the human UniprotID, position number of the phosphorylated residue in the protein sequence, and the log2 FC of the phosphorylation site of a given treatment.

